# Development of a cellular reporter assay to measure activity of MutSβ, a therapeutic target for Huntington’s disease

**DOI:** 10.1101/2023.09.07.555786

**Authors:** Jian An, Theresa Towle, Melis Atalar Aksit, Mohiuddin Mohiuddin, Samantha Castaneda, Reiko Nakashima, Rob Moccia, Christine Bulawa, James Fleming

**Author notes:** Correspondence: Christine Bulawa, PhD, and Jian An, PhD, Rare Disease Research Unit, Worldwide Research, Development and Medical, Pfizer Inc., 610 Main Street, Cambridge, MA, 02139, USA. C.B. and J.F. contributed equally to this manuscript.

## Abstract

Genetic modifiers of age of onset in Huntington’s disease (HD) provide compelling evidence that somatic expansion of the CAG repeats is a critical driver of pathogenesis and demonstrate that repeat instability is modulated by DNA mismatch repair (MMR). A component of this pathway, MutSβ, a heterodimer comprised of MSH2 and MSH3, has emerged as a potential target for small-molecule therapeutic intervention. However, a robust cellular assay to interrogate genetic and pharmacological modifiers of MutSβ has not been reported. We have repurposed and optimized a tetranucleotide reporter assay to measure MutSβ activity in MMR-competent cells. We show that repeat instability is modulated by MSH3 protein levels and by its ATPase activity. In addition, we show that an inhibitor of HDAC3 modulates repeat instability, demonstrating the utility of the assay for pharmacological studies.

## Introduction

Microsatellites are short tandem DNA repeats found throughout the human genome. The expansion of these repetitive sequences underlies >40 human diseases including Huntington’s disease (HD), C9orf72 ALS/FTD, myotonic dystrophy, and spinocerebellar ataxias (Depienne and Mandel 2021; Paulson 2018). These repeat expansion diseases (REDs) are severe, progressive disorders for which there are no disease modifying therapies available to patients. Across the REDs, the monomer repeat length, sequence, and position within the associated locus can vary; however, several commonalities are observed including: 1) An inverse correlation between repeat length and disease age of onset and severity. 2) A pronounced increase in the expansion of repeats in germline which is responsible for anticipation (the decreased age at onset and increased disease severity in successive generations). 3) Somatic expansion of the repeats, that is an increase in the repeat length above the inherited number, which occurs over the lifetime of the individual, attaining sizes of 100’s to 1000’s of repeats (Kennedy et al. 2003). Strikingly, expansion is most severe in disease affected tissues, especially in non-dividing cell types such as neurons. Moreover, the degree of somatic instability is associated with age of onset, and interruptions of the repeats that decrease somatic instability are associated with later disease onset (Wright et al. 2019; Genetic Modifiers of Huntington’s Disease Consortium. Electronic address and Genetic Modifiers of Huntington’s Disease 2019). These data support repeat instability as a driver of disease pathogenesis.

Human genome-wide association studies have been performed to identify factors (Genetic Modifiers of Huntington’s Disease 2015) in addition to repeat length that contribute to age of onset and disease progression in HD. Results of these studies support a two-step model for pathogenesis in which the initial step is the expansion of an inherited disease allele and the subsequent step is cellular dysfunction/loss that occurs when repeats exceed a threshold size (Hong et al. 2021). Notably, these studies identified multiple genes involved in DNA mismatch repair (MMR) and indicated that loss of MMR function correlates with delays in age of onset and lowered disease severity. Additionally, individual knockout of several components of the MMR pathway in mouse models of HD and other REDs reduces repeat expansion (Wheeler and Dion 2021). Therefore, proteins in this pathway serve as potential drug targets for the REDs by blocking the first step in the two-step model of pathogenesis (Benn, Gibson, and Reynolds 2021).

There are 2 branches of the MMR pathway with a number of overlapping protein components (Iyer and Pluciennik 2021). These branches can be distinguished by the sentinel protein complexes, MutSα and MutSβ, that initiate mismatch repair by detecting DNA lesions and recruiting additional factors that catalyze repair. MutSα (composed of MSH2 and MSH6) identifies and initiates repair of small loop outs of 2 or less nucleotides as well as mis-paired bases, while MutSβ (composed of MSH2 and MSH3) detects and repairs larger (≥2 nucleotide) loop outs. MSH3, the unique component of the MutSβ complex, was first identified as a major determinant of somatic instability in mice (Dragileva et al. 2009) and was subsequently found to modify age of onset and disease progression in HD patients. Other MMR proteins also modulate somatic instability, but from the perspective of drug development, MSH3 has two distinct advantages. First, repeat instability is very sensitive to the level of MSH3. In mice, heterozygous KO of MSH3 significantly inhibits expansion whereas no effect is seen in heterozygous knockouts of other MMR modifier genes (Dragileva et al. 2009). Second, MSH3 inhibition carries minimal cancer risk (Aelvoet et al. 2023; Adam et al. 2016). Therefore, MSH3 has emerged as a potential therapeutic target for HD and potentially additional REDs.

To facilitate the discovery and development of modulators of MSH3, we have established a robust cell-based assay of repeat instability. We show that a mutation in the ATPase domain fails to complement an MSH3 knockout, supporting the ATPase domain as a therapeutic target. In addition, we show that an inhibitor of HDAC3 modulates repeat instability, demonstrating the utility of the assay for pharmacological studies.

## Results

### AAAG frame shift reporter cell line generation and characterization

To study modulators of MSH3, an effective cell-based assay of repeat instability is needed. Major challenges to assay development include the low frequency of instability events and the lack of facile methods to measure repeat length. Most published cellular assays of CAG instability require 4-8 weeks in culture (Goold et al. 2019; Roy et al. 2021; Nakamori et al. 2020), and the throughput of methods to determine repeat length is inadequate for screening large libraries. For long repeats, >100 CAG, the observed expansion is typically only 3-5 repeats, which limits the sensitivity and robustness of this approach.

An alternative to the CAG instability assay is based on the role of MSH3 in Elevated Microsatellite Alterations at Selected Tetranucleotide repeats (EMAST), a type of microsatellite instability (MSI) observed in multiple cancer types, especially colorectal cancer (CRC). Tetranucleotide repeats, in contrast to trinucleotide repeats, become more unstable when MSH3 function is reduced. Campregher et al. (Campregher et al. 2012) developed a cellular EMAST assay based on the instability of AAAG, a major tetranucleotide repeat in the human genome. These investigators created cells expressing a reporter gene with 17 concatenated AAAG repeats fused to green fluorescent protein (GFP) (Figure 1A). Cells with this 17-repeat reporter were GFP negative (GFP-) because GFP is not in frame with the initiating ATG. Over time a subpopulation of GFP positive (GFP+) cells were generated. FACS sorting and DNA sequencing demonstrated that the number of repeats in the GFP+ cells was changed from 17 to a multiple of 3, thereby bringing GFP in-frame with the initiating ATG. Knock-down of MSH3 enhanced this instability, consistent with results from EMAST studies in cancer patients.

**Figure 1.**
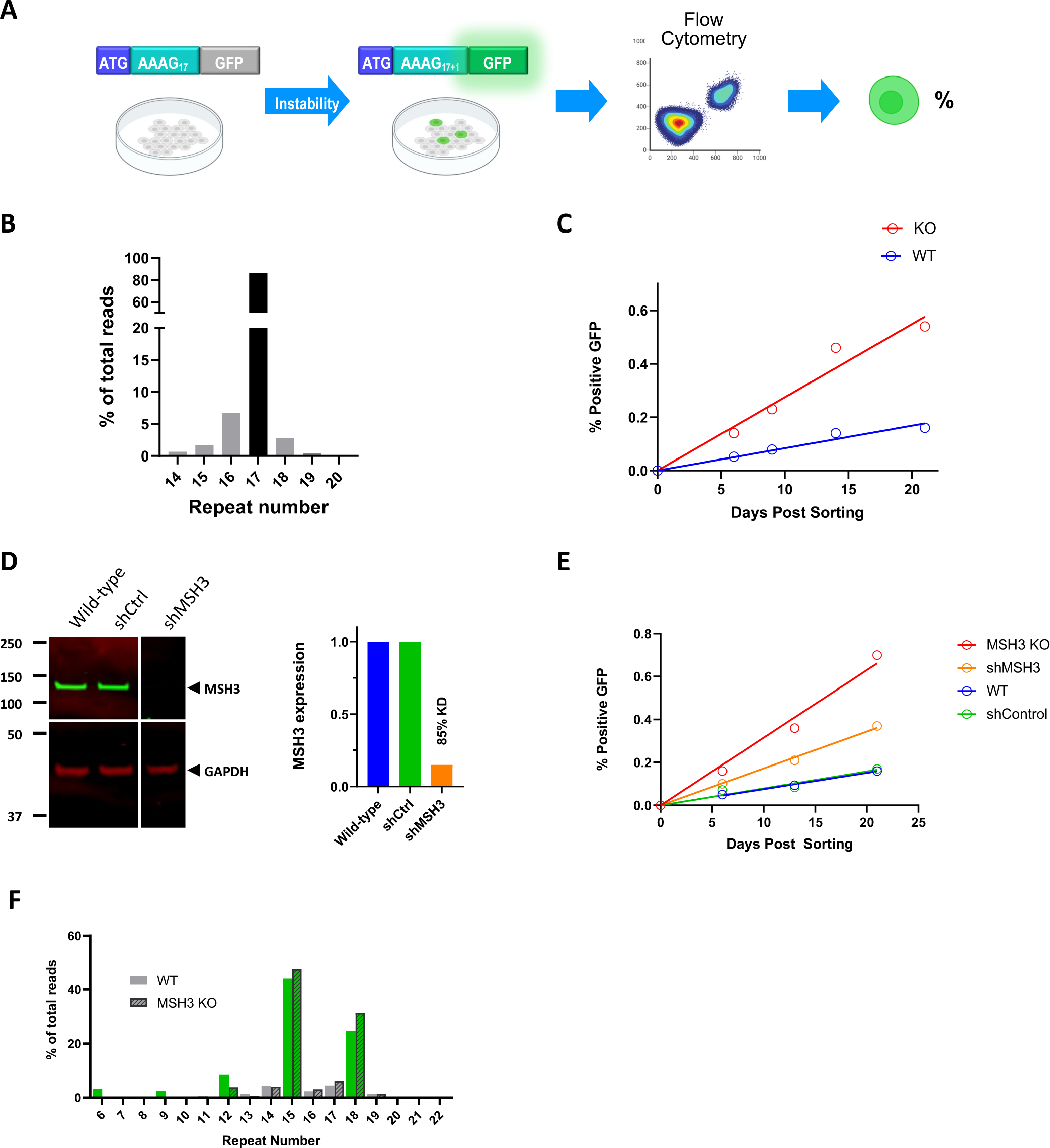
Characterization of AAAG_17_ reporter line. A. Cartoon illustrating frameshift assay concept. B. Distribution of repeat counts in RNA-seq reads from AAAG_17_ reporter cell line; expressed as percent of total reads. Starting length is indicated by dark bar. For visualization purposes, repeats with 1-3 base additions or deletions (0.6% of all reads) were excluded from the plot. C. Kinetics of GFP+ conversion in Flp293 wild-type (blue) and MSH3 KO (red) AAAG_17_ reporter lines (n=1). D. Western blot probing MSH3 protein levels in wild-type cells, cells stably expressing shRNA directed toward MSH3, and cells stably expressing control shRNA (left). Quantitation of MSH3 knockdown effect normalized to GAPDH (right) (n=1). E. Kinetics of GFP+ conversion in wild-type (blue), MSH3 KO (red), control shRNA (green), and MSH3 shRNA (orange) AAAG_17_ reporter lines. (n=1). F. Distribution of repeat count in RNA-seq reads from GFP+ cells from wild-type (solid bars – left) and MSH3 KO (hatched bars – right) AAAG_17_ reporter cells lines; expressed as percent of total reads. Green color represents repeat numbers that would produce in frame GFP protein expression. Repeats with 1-3 base additions or deletions were excluded from the plot (1.2% and 0.18% of reads in the wild-type and MSH3 KO cells, respectively).

The EMAST assay has certain advantages compared to published CAG based assays. The increase in tetranucleotide instability, although directionally opposite to trinucleotide repeat instability, has the benefit of increasing the fluorescence signal in response to MSH3 knockdown and utilizes flow cytometry, which is amenable to miniaturization and automation. We therefore set out to generate an improved assay with increased robustness to enable both mutational analyses and compound screening. Our frameshift assay utilizes an HEK cell line, Flp-In™ 293, commonly employed in cell-based screening. This line contains a locus for the insertion of the reporter constructs, which limits clonal variation in expression arising from differences in the site and/or number of integration events. Because many cell lines harbor perturbations of the MMR pathway, we confirmed the expression level (Figure S1) and found no predicted loss-of-function or ClinVar pathogenic variants (Table S1) in the known components of the MMR pathway in the Flp-In™ 293 cells, indicating that the MMR pathway is expected to be functional in this cell line. We then generated lines expressing a single copy of the GFP transgene with 17 repeats inserted between the ATG and the triplet coding for amino acid 2 in GFP in both wild-type and MSH3 knockout cells. A wild-type line with an in-frame GFP reporter (18 repeats) was also generated to confirm that the peptide sequence encoded by the repeat sequence did not significantly impact GFP fluorescence and to serve as a positive control for the flow gating strategy. After clonal selection, droplet digital PCR confirmed the integration of a single copy of the transgene inserted at the flippase recognition target (FRT) site (Figure S2), and the number of the repeats in the reporter was confirmed via RNA-seq (Figure 1B).

To establish the rate of GFP+ conversion and its dependency on MSH3, we sorted wild-type and MSH3 knockout clones to obtain pure GFP-populations and monitored the generation of GFP positive cells over the course of three weeks. In both cell lines, the rate of GFP positive cells was constant over the course of the assay. In the wild type cells ∼0.03% of cells converted to GFP+ per week. This rate was increased 5-fold in MSH3 knockout cells, to 0.16% per week (Figure 1C). To ensure that this increase was due to the deletion of MSH3, we re-introduced WT MSH3 using a lentivirus vector, which restored the low GFP+ conversion rate observed in the WT line (Figure 3A). Further testing demonstrated the assay displays excellent intra-experiment, interexperiment and clonal reproducibility (Figure S4) with an increase in the GFP+ conversion rate in MSH3 null cells relative to wild-type cells that ranged from 3- to 5-fold across experiments.

To assess the effect of a partial loss of MSH3 function in the HEK cells, we generated stable cell lines expressing either a scrambled shRNA or a shRNA targeting MSH3 which reduced protein levels by approximately 85% in the AAAG_17_ reporter cell line (Figure 1D). Cells expressing the scramble shRNA control had an equivalent rate of GFP+ conversion as wild-type cells, while the cells with 15% residual MSH3 protein levels had a GFP conversion rate intermediate between the wild-type and MSH3 knockout clones (0.1% per week, Figure 1E). Of note, the change in GFP+ conversion rate is not proportional to the change in MSH3 protein level. In cells with 85% MSH3 knockdown, the GFP+ conversion was 60% lower than the rate of observed in the MSH3 KO cells, in contrast to the expected 15% reduction. The reason for this limited effect is unclear (see discussion).

To determine whether the gain of GFP signal in the assay is due to change in repeat number or whether it occurs by a different mechanism, (e.g., 1 nucleotide insertion or 2 nucleotide deletion), we sorted wild-type and MSH3 KO cells into GFP+ populations at the end of a 21-day experiment and performed RNA-seq. Analysis of reads that spanned the AAAG repeat sequence demonstrates that the majority of reads (> 83%) had a repeat number that generated an in-frame GFP reporter. A large proportion of the in-frame reads consisted of 15 or 18 repeats suggesting the unit change in repeat length is typically small (Figure 1F); 1-3 base additions or deletions were rarely detected in the repeats (1.2% and 0.18% of reads in the wild-type and MSH3 KO cells respectively, Table S2). A similar analysis of a population sorted for GFP-cells demonstrated that ∼98% of reads contained repeat numbers that would lead to an out-of-frame GFP reporter as expected (Figure S5). These data demonstrate the suitability of using the GFP+ conversion rate as a surrogate readout for instability of the AAAG repeat.

### Optimization of the frameshift assay

While the AAAG_17_ assay reproducibly monitors repeat instability, its utility is limited by the long incubation time required for an adequate signal. Analysis of our RNA-seq data provided a potential avenue for improvement. RNA sequencing of the unsorted AAAG_17_ cells reveals that the predominant repeat unit change is one repeat, and contractions of a single repeat occur approximately 2.5 times more frequently than expansions (Figure 1B). Therefore, we hypothesized that a reporter with a single additional repeat unit above an in-frame number would result in an increase in the GFP conversion rate, even if the underlying instability rate remains constant. We generated additional cell lines with reporters containing 13, 19 and 31 AAAG repeats upstream of GFP and evaluated these in the assay. The GFP+ conversion rate in wild-type cells was greatest for the AAAG_31_ line and lowest for the AAAG_13_ line (Figure 2A) as expected based on the correlation of instability to repeat length. Because we ultimately wanted an assay with a shorter incubation time, we characterized the AAAG_31_ reporter further. We assessed the dependence of GFP+ conversion on MSH3 using the knockdown and knockout approaches described above for the AAAG_17_ reporter. For each approach, we purified and characterized two independent clones. The two strains expressing shRNA directed at MSH3 had low residual levels of MSH3 protein as expected (Figure S6, panel C); however, the increase in GFP+ conversion rate was small relative to shRNA controls and WT (Figure S6, panel A) and therefore did not provide an advantage relative to the AAAG_17_ reporter. Knockout of MSH3 in the AAAG_31_ strains produced a greater effect but unexpectedly, re-introduction of MSH3 failed to fully complement the instability (Figure S6, panel B). Analysis of AAAG_19_ reporter lines revealed significant clonal variation (data not shown) thus precluding further study with this line. In conclusion, characterization of the AAAG_19_ and AAAG_31_ lines revealed technical challenges and unanticipated results, and they were not pursued further.

**Figure 2.**
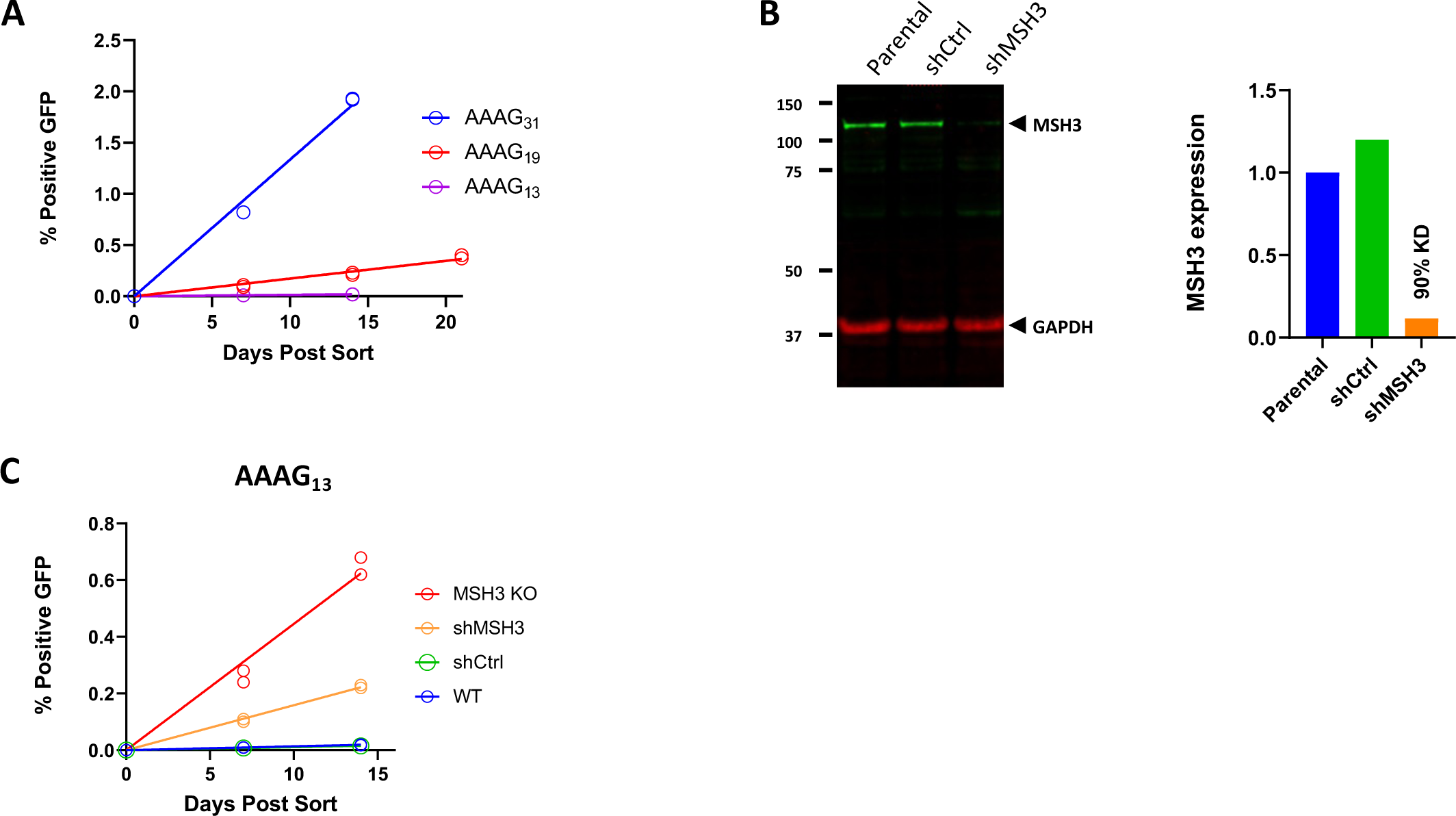
Characterization of additional AAAG repeat length reporter constructs. A. Kinetics of GFP+ conversion in reporter lines with 31 (blue), 19 (red), and 13 (purple) AAAG repeats. Replicate (n=2) data points are plotted (in some instances replicates are indistinguishable). B. Western blot probing MSH3 protein levels in AAAG_13_ wild-type cells, cells stably expressing shRNA directed toward MSH3, and cells stably expressing control shRNA (left). Quantitation of MSH3 knockdown effect normalized to GAPDH loading control (right) (n=1). C. Kinetics of GFP+ conversion in wild-type (blue), MSH3 KO (red), control shRNA (green), and MSH3 shRNA (orange) AAAG_13_ reporter lines. Replicate (n=2) data points are plotted (in some instances replicates are indistinguishable).

In contrast to the AAAG_31_ and AAAG_19_ reporters, AAAG_13_ displayed favorable characteristics, performing as well or better than AAAG_17_. As described above, in MSH3 WT cells, AAAG_13_ instability was low relative to AAAG_17_. In MSH3 null cells, the GFP+ conversion rate for AAAG_13_ (Figure 2C, 0.3 repeats/week) was nearly 1.5- to 2-fold greater compared to AAAG_17_ (Figure 1C, E, 0.15 - 0.2 repeats/week). As anticipated, GFP+ cells arise primarily by loss of one repeat (Table S3). In both strains, shRNA mediated knockdown of MSH3 (∼90% reduction, Figures 1D and 2B) led to a similar increase in GFP+ conversion rate relative to MSH3 KO cells (Figures 1E and 2C). In addition, we observed stronger fluorescence for the AAAG_13_ reporter, which provides better separation between the GFP+ and GFP-cell populations (Figure S7). Based on these data, the AAAG_13_ and AAAG_17_ lines are both suitable for monitoring MSH3 activity and were used to evaluate the effects of modulation of MSH3 on repeat instability as described in the studies below.

### MSH3 ATPase activity is required for its effect on AAAG stability

MSH3 belongs to the ABC ATPase family and contains Walker A and Walker B motifs, which are responsible for nucleotide binding and ATP hydrolysis. In vitro and in vivo studies have demonstrated that lowering the level of MutSβ protein halts repeat expansion, but the dependence of repeat expansion on the ATPase activity of MutSβ has not been unequivocally established (Keogh et al. 2017; Edwards 2017). To address this question, we tested the ability of the Walker B mutant, E976A MSH3 to complement the MSH3 KO. We first confirmed published results (Keogh et al. 2017) demonstrating the lack of ATPase activity in recombinant MutSβ with an E976A mutation in the MSH3 subunit (data not shown). Then, AAAG_13_ and AAAG_17_ MSH3 KO cells were stably transduced with lentiviral vectors carrying either MSH3 WT or the E976A mutant and assayed to determine the rate of generation of GFP+ cells.

While complementation with wild-type MSH3 reduced the frameshift rate to the level seen in parental (WT) cells, the E967A mutant complementation had no effect on the frameshift rate in either the AAAG_13_ or AAAG_17_ MSH3 KO strains (Figure 3A). Importantly, we observed that the E976A mutation did not alter the protein level in either strain (Figure 3B and S8), the nuclear localization of MSH3 protein (nuclear fractionation, Figure 3C) or its association with MSH2 (co-immunoprecipitation, Figure 3B) in the E976A complemented AAAG_17_ cells (Figure 3B). Because our knockdown data shows that even a low residual amount of MSH3 protein significantly stabilizes the repeats (the instability in a strain with 10-15% residual MSH3 protein level is 35-40% relative to the MSH3 KO GFP+ rate defined as 100%), we infer that the residual cellular activity of the E976A mutant is likely very low, <10% of WT.

**Figure 3.**
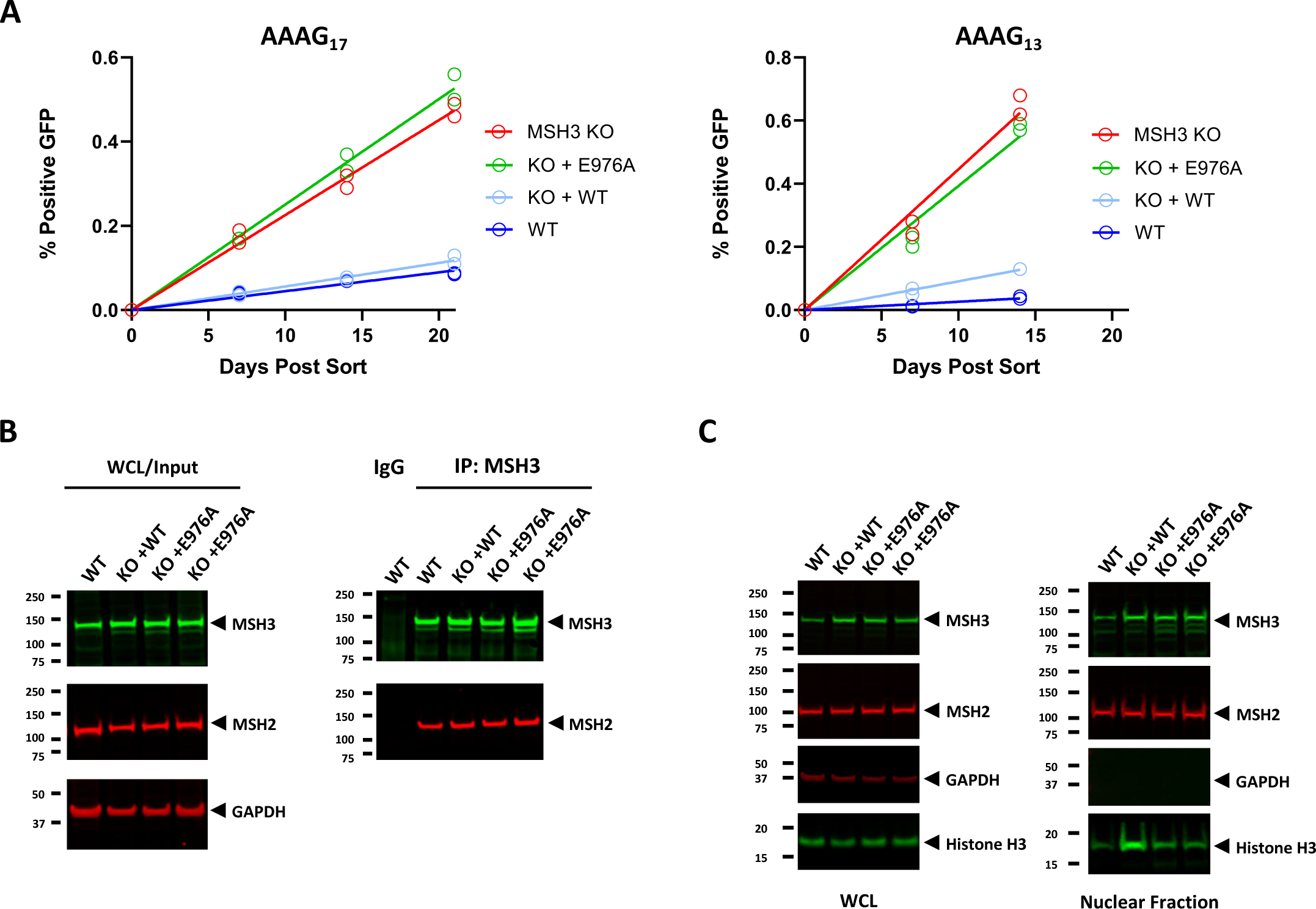
Genetic complementation. A. Kinetics of GFP+ conversion in wild-type (dark blue), MSH3 KO (red), MSH3 KO cells expressing wild-type MSH3 (light blue), and MSH3 KO cells expressing the E976A MSH3 mutant (green) in AAAG_17_ (left) and AAAG_13_ (right) reporter lines. Replicate (n=2) data points are plotted (in some instances replicates are indistinguishable). B. Western blot probing MSH2 and MSH3 protein levels in the AAAG_17_ reporter lines from whole cell lysates (WCL - left) and from samples after immunoprecipitation with a MSH3 directed antibody (right). C. Western blots probing MSH2, MSH3, GAPDH and Histone H3 protein levels in whole cell lysates (WCL - left) and in nuclear extracts from the indicated AAAG_17_ reporter lines.

### Known coding polymorphisms do not alter MSH3 function in frameshift assay

Two MSH3 variants of high interest, designated 3a and 7a, are associated with HD age of onset. Both reside in exon 1 in a polymorphic region containing 9 bp tandem repeats (Flower et al. 2019). Compared to the major allele, 6a, the 3a allele has a shorter alanine repeat, and the 7a allele has three additional amino acids, PAA (Figure S9A). The effect of these variants on MSH3 function is not well characterized. In the case of 3a, expression quantitative trait loci data suggest a modest reduction of MSH3 level (Flower et al. 2019) which is unlikely to be detectable in the frameshift assay because high levels of MSH3 knockdown are required to alter the rate of GFP+ conversion. Nonetheless, because of their disease relevance, we generated stable lines expressing each individual allele in AAAG_17_ MSH3 KO cells and measured the frameshifting rate. As expected, no difference was detected between the 6a, 3a, and 7a polymorphisms (Figure S9).

### Testing small molecules in frameshift assay

To ascertain whether the frameshift assay would be amenable for compound screening, we evaluated the HDAC3 inhibitor RGFP966. Previous studies indicate that histone deacetylases (HDAC) promoted the instability of CAG repeats, and chronic administration of RGFP966 in cells or in the Q111 mouse model of HD suppressed CAG repeat expansion (Suelves et al. 2017) potentially through a direct effect on MutSβ (Williams et al. 2020). We chronically treated the AAAG_13_ and AAAG_17_ reporter cells with RGFP966 and observed an increase in the rate of GFP+ conversion (Figure 4). The rate of GFP+ conversion in the RGFP966 treated cells was 1.9-fold greater in the AAAG_17_ cells and 3.6-fold greater in the AAAG_13_ line when compared to their respective DMSO controls, effect sizes that were small relative to the MSH3 KO controls (14% and 8% for AAAG_17_ and AAAG_13_ respectively). Importantly for screening purposes, the RGFP966 response was statistically significant at 2 weeks for both reporter strains and at 1 week for the AAAG_13_ reporter strain even with the limited number of replicates employed and small effect size observed.

**Figure 4.**
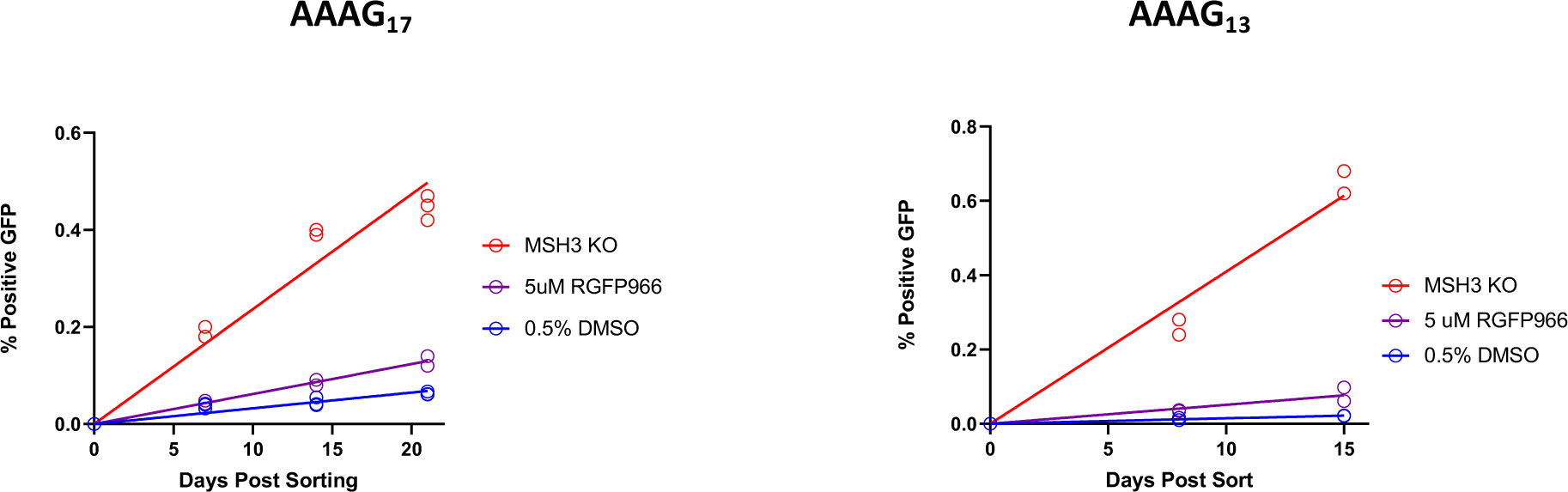
RGFP966 modulation of GFP+ conversion rate. Kinetics of GFP+ conversion in MSH3 KO (red), DMSO treated wild-type (blue) and RGFP treated wild-type AAAG_17_ and AAAG_13_ reporter lines. Replicate (AAAG_17_ n=3, AAAG_13_ n=2) data points are plotted (in some instances replicates are indistinguishable).

## Discussion

We have developed and characterized a robust cellular assay to monitor MutSβ activity. This work repurposes and improves the assay described in Campregher et al. (Campregher et al. 2012) who used an AAAG reporter to explore the relationship between EMAST and carcinogenesis. Using MSH3-deficient cancer cell lines and silencing of MSH3 in primary colonic epithelial cells, they concluded that loss of MSH3 drives EMAST but not oncogenic transformation. We have modified the tetranucleotide reporter described by Campregher et al. (Campregher et al. 2012) and created MMR-proficient cell lines that robustly monitor genetic and pharmacological modulation of MSH3, a target of interest for HD and other REDs.

We evaluated multiple parameters during the development of our frameshift assay. We wanted an MMR-proficient cell line that allows robust comparison of different reporters and genetic perturbations. We chose the well-characterized Flp-In™ 293 cell line to integrate the AAAG reporters into the genome to control for both copy number as well as genome integration site. Using the AAAG_17_ construct, we first confirmed that instability is dependent on MSH3 and then evaluated additional repeat lengths to identify the optimal number for assay sensitivity and duration. AAAG_19_ and AAAG_31_ constructs have increased rates of instability as anticipated (Figure 2A); however, the unexpected behavior and clonal variability of these lines in knockdown and complementation experiments led to their discontinuation. In contrast, the AAAG_13_ reporter line has multiple characteristics that support its utility for screening. First, the AAAG_13_ GFP+ cells had the highest degree of fluorescence and therefore the greatest window for gating between the GFP- and GFP+ populations (Figure S7). The stronger fluorescence of the AAAG_13_ reporter is consistent with our observation of an inverse relationship between AAAG length and GFP fluorescence signal. This property is particularly desirable for drug screening as compounds can interfere with signal detection either directly by absorbing/fluorescing in the GFP excitation/emission spectra, or indirectly by increasing cellular autofluorescence. Second, while the rate of GFP+ conversion in wild-type cells was lower for the AAAG_13_ vs AAAG_17_ line as expected, the rate of GFP+ conversion in MSH3 KO cells was greater for the AAAG_13_ reporter lines (Figures 1C vs 2C and Figure 3). This improves the assay window for the AAAG_13_ line, providing a readout in a shorter time.

We developed the frameshift assay to support the discovery and optimization of small molecule inhibitors of MutSβ, but its utility is much broader. The frameshift assay is likely applicable to other enzymes associated with microsatellite instability, such as MSH2, MLH1, and PMS2 (Raeker and Carethers 2020), a possibility that can be readily interrogated by gene knockout experiments. The resulting strains can then be used for complementation studies to explore structure-function relationships as we have done for the E976A variant of MSH3. Similar repeat instability assays exist for yeast (Sia et al. 1997) and have provided mechanistic insights into variants of MSH3 (Kumar et al. 2013). Finally, the frameshift assay is well suited to discover new modifiers of repeat instability by performing genome-wide screens to identify genes whose loss of function or overexpression modulate the rate of formation of GFP+ cells and therefore repeat instability.

The HDAC inhibitor RGFP966 has been reported to inhibit expansion of CAG repeats in Q111 knock-in mice. A selective inhibitor of histone deacetylase 3 prevents cognitive deficits and suppresses striatal repeat expansions in Huntington’s disease mice (Suelves et al. 2017) and in cells (Williams et al. 2020). We show that RGFP966 also has modest activity in the frameshift assay, but testing at higher concentration was not possible due to cytotoxicity. It has been suggested that acetylation regulates the distribution of MSH3 between the nucleus and cytoplasm and enhances localization to the cytoplasm. This hypothesis, however, has not been demonstrated unequivocally (M.E. Herva, CHDI 18th Annual HD Therapeutics Conference, April 24-27, 2023, Abstract 92). Despite the ambiguity of its molecular mechanism, the effect of RGFP966 was reproducible in our hands, and we believe that it can serve as a positive control.

The assay described herein robustly monitors instability of the AAAG repeat. However, there are limitations. First, while we have shown that modulation of instability is detectable in as little as 1 week for the AAAG_13_ construct and 2 weeks for the AAAG_17_ construct, accelerating the rate further is desirable. We optimized AAAG length, but there are many additional parameters that could be explored. For example, it is possible that a different repeat may display greater instability which should shorten the assay duration. Second, our current assay requires large numbers of cells (>100,000) for robust determination of the frameshift rate. Further modifications, such as use of a more sensitive reporter (e.g., luciferase) may require fewer cells, thereby allowing miniaturization and/or automation. Third, our data show that a large degree of knockdown of MSH3 protein level (85-90%) only produces a modest effect on instability relative to a complete MSH3 null. This is inconsistent with in vivo data demonstrating that animals expressing a single functional copy of MSH3 (MSH3+/null) have a profound lowering of repeat instability in the striatum (Dragileva et al. 2009). Thus, sensitizing our assay to MSH3 inhibition is desirable. This might be possible by using a different cell line or by lowering MSH3 levels, for example by knocking out one allele or using siRNA. Finally, in the AAAG tetranucleotide repeat assay the loss of MSH3 function increases instability. To confirm their relevance to REDs, genetic and/or pharmacological perturbations that modulate instability in this system will need to be evaluated for their effect on instability of CAG repeats for which loss of MMR function decreases instability.

We note that similar assays have been reported. Raeker et al, (Raeker, Pierre-Charles, and Carethers 2020) designed reporters containing the human D9S242 tetranucleotide microsatellite locus and characterized EMAST in MMR-deficient HCT116 cells. During the preparation of this manuscript, several new repeat instability assays have been presented. Recently, at the CHDI 18th Annual HD Therapeutics Conference, April 24-27, 2023, three groups presented AAAG assays designed to monitor MSH3 activity (abstract 50, Sara Tomaselli, IRBM, abstract 88, Karsten Tillack, Evotec SE, and abstract 93, Michael D Flower, University College London) and McLean et al. published a preprint for a new assay for CAG instability (https://www.biorxiv.org/content/10.1101/2023.07.25.550489v1).

In summary, we have developed a cellular frameshift assay that can reliably monitor AAAG repeat instability. We demonstrate that our assay can identify genetic and pharmacological modulators of MSH3, a modifier of HD age of onset, and that it is likely applicable to additional components of DNA MMR.

## Methods

### Generation of the pCDNA5 FRT-[AAAG]17 or 13, 19, 31-EGFP-frameshift reporter plasmid

EGFP (Sequence ID: MN443913.1) reporter constructs flanked by BamHI and EcoRV restriction sites were synthesized (GeneWiz) with varying numbers of AAAG repeats inserted between the ATG start codon and the codon for the second amino acid. These sites were then utilized to clone the reporter construct into the pCDNA5 FRT multi-cloning site (ThermoFisher, V601020).

### Generation of AAAG_(n)_-eGFP Flp-In™ 293 cell lines

The Flp-In™ 293 cell line was obtained from ThermoFisher (R75007). Cells were maintained in DMEM (Gibco, 11995-065) supplemented with 15% fetal bovine serum (FBS; Gibco, 16000044), 20mM HEPES buffer (Gibco, 15630-080), 20 U/mL penicillin, and 20 µg/mL penicillin/streptomycin (Gibco Pen/Strep, 15140-122). Cell lines were generated as per the manufacturer’s protocol. The day prior to transfection 10 cm dishes were seeded with 2× 10^6^ cells. Cells were transfected using a master mix of Fugene 6 (Promega, E269A) and Optimem (Gibco, 31985-070) containing 9 µg pOG44 and 1 µg of the reporter construct plasmid and incubated for 15 minutes. 1 mL of transfection complex was then added to a corresponding 10 cm dish of Flp-In™ 293 cells. Medium was replaced approximately ∼24 hours following transfection. Medium containing 150 µg/mL hygromycin (Invitrogen, 10687010) was exchanged ∼48h post-transfection. Clones were selected via limiting dilution under selection conditions. After clonal isolation cells were maintained in 150 µg/mL hygromycin.

### Generation of human MSH3 knockout cell lines

Oligonucleotide sequence ‘5-AGAGACCATTGGAAAATGAT-3’ was cloned into pX330-U6-Chimeric_BB-CBh-hSpCas9 plasmid with a fluorescent reporter gene. The vector was transfected into Flp-In™ 293 cells using Fugene 6 (Promega, E269A) and 24 hours later the cells expressing the fluorescent reporter were sorted in a 96 well plate as single cells using the Sony SH800 cell sorter. MSH3 knockout was confirmed by Western blot.

### Generation of knock down AAAG13 cell lines

24 hours prior to transduction 0.2 million cells/well were seeded in a 6-well plates. The next day, media was exchanged with media containing 10 µg/mL polybrene (EMD Millipore, TR-1003-G) and either shControl (pLKO.1-puro Sigma Aldrich) or shMSH3#59 (TRCN0000417959 Sigma Aldrich) lentivirus at MOIs of 2. Medium containing 2 µg/mL puromycin (Gibco, A11138-03) was exchanged 24 hours post-transduction. Cells were grown under 2 µg/mL puromycin selection for one week followed by maintenance in 0.5 µg/mL puroymycin.

### Generation of complemented AAAG13 cell lines

24 hours prior to transduction, 2 million cells were seeded in 10 cm dishes. The next day, media was exchanged with media containing 10 µg/mL polybrene (EMD Millipore, TR-1003-G) and lentivirus containing a cassette that will express either wild-type MSH3 or an E976A mutant MSH3 (triplet change from GAA to GCA) at MOIs of 5. Medium containing 2 µg/mL puromycin (Gibco, A11138-03) was exchanged 24 hours post-transduction. Cells were grown under 2 µg/mL puromycin selection for one week followed by maintenance in 0.5 µg/mL puromycin.

### Evaluation of AAAG reporter copy number in Flp-In™ 293 cell lines

Cells from 1 well of a 96 well plate were used as samples for gDNA extraction. 80 µL of Lucigens QuickExtract DNA Extraction Solution (QE0905T) was used following the manufacturer’s protocol. 50 ng DNA input was run in BioRad digital droplet PCR (ddPCR™) using dUTP-free ddPCR Supermix for Probes (BioRad, 1863024). Reactions were set up according to manufacturer’s instructions, with droplets prepared using the Auto DG droplet generator (Bio-Rad). Amplification of target genomic DNA was performed using the following cycling conditions: 95°C for 10 minutes x 1, 94°C for 30 s x 40, 60°C for 1 minute x 40, 98°C for 10 minutes x 1. Droplets were then read using a QX200 Droplet Reader (Bio-Rad) and data were analyzed using Quantasoft software (Bio-Rad) following manufacturer’s recommendations for quality control and analysis. A custom GFP primer/probe set (Forward sequence: AGCAGAAGAACGGCATCAA,Hairpin BlastProbe: CAAGATCCGCCACAACATCGAGGA, BlastReverse sequence: GTGCTCAGGTAGTGGTTGTC, label:FAM) was employed, and Tert was used as the reference gene (VIC labeled: ThermoFisher, 4403316).

### Co-Immunoprecipitation

24 hours prior to transduction, 4 million cells were seeded in 10cm dishes. The next day cells were washed with PBS followed by dissociation with trypsin (ThermoFisher Scientific 25200-056). Cell pellets were obtained by centrifugation at 300 xg for 5 minutes and washed with PBS two times. Lysis was performed by the addition of 1mL IP lysis buffer (150mM NaCl, 50mM Tris pH 7.5, EDTA 1mM, 0.5% NP-40 and 10% glycerol) containing protease inhibitor cocktail (cOmplete Roche, 04693159001), followed by end-over-end rotation at 4 °C for 6 hours. Lysates were clarified by centrifugation at 16,000 xg for 30 minutes and the supernatant was added to 20 µL of pre-washed Protein G bead slurry (ThermoFisher Scientific, 20398). The following antibodies were added to their respective tubes, 10 µL anti-MSH3 antibody (Bethyl Labs anti-MSH3 A305-314A) and 5 µL Control Rabbit IgG (Invitrogen, 10500C), then incubated with end-over-end rotation at 4 °C overnight. Tubes were then spun down at 200 xg for 5 mins and the supernatant was removed. Beads were washed with 1mL IP lysis buffer followed by centrifugation at 200 xg for 5 minutes, six times. Western blot buffer 4X NuPAGE LDS sample buffer (Invitrogen, NP0007) and 10X NuPAGE Sample Reducing Agent (Invitrogen, NP0004) diluted in IP buffer was added to the beads. Samples were heated at 95 °C for 5 minutes and used for Western blot analysis.

### RNA-seq

Cell pellets were sent to Fulgent for processing. The RNA concentration was determined using QuBit RNA High Sensitivity Kit from ThermoFisher (Q10211, Q32855). RIN measurements were determined by using the 4200 Tapestation System from Agilent (5067-5579). 500 ng of RNA was used as an input and libraries were prepared using the TruSeq Stranded mRNA library preparation kit from Illumina. The samples were sequenced on the NovaSeq 600 platform on the NovaSeq S4 300 cycle flowcell to generate 150 bp paired-end reads. Global alignment against the plasmid flanking sequences that are located 60 bp upstream and 62 bp downstream of the AAAG repeat sequence was performed with BBMap version 38.21. The reads that span the entire AAAG repeat region were extracted with Jvarkit version 2021.10.13 after reformatting with SAMtools, version 1.13. The number of AAAG repeats was determined with a custom Python script (version 3.8.16) by counting the bases in the repeat region, starting from the first AAAG + 4bp upstream flanking sequence through the last AAAG + 4bp downstream flanking sequence. The repeat length was calculated as the total number of bases between the first and last AAAG, and the repeat count was estimated as the repeat length divided by four. For visualization purposes, the reads with a repeat length that were not a multiple of four (0.1-1.2% of total reads) and some repeat lengths with low representation (< 0.5% of total reads) were excluded from the plots. (Supplemental Tables 1 and 2 contain the full data sets.)

### Western blotting

Whole cell lysates were collected from 1 well of a 12 well plate using RIPA lysis buffer (1% NP-40, 0.5% sodium deoxycholate, 200mM NaCl, 20mM Tris-HCL pH7.5, 1mM EDTA) containing HALT protease/phosphatase inhibitors (ThermoFisher 78430). Samples were centrifuged for 30 minutes at 4 °C and the supernatant was transferred to a new tube. Protein concentration was determined using Pierce 660nm Protein Assay Kit (ThermoFisher, 22662). 12 µg protein plus 4X NuPAGE LDS sample buffer (Invitrogen, NP0007) and NuPAGE Sample Reducing Agent (10x, Invitrogen, NP0004) were added. After heating at 95 °C for 5 minutes, samples were loaded on the gel. SDS-PAGE was performed using a 4-12% gradient gel (Invitrogen, WG1402BOX) and MES running buffer (ThermoFisher, NP0002). Proteins were transferred to nitrocellulose membranes (Bio-Rad, Turbo Midi trans-blot, 1704159). Membranes were blocked for 1 hour at room temperature (LI-COR Intercept TBS blocking buffer, 927-60001) and were probed overnight at 4 °C with primary antibody (see table below). Membranes were washed and incubated with anti-rabbit IgG (LI-COR, IRDye® 800CW, 926-32211, 1:20,000) or anti-mouse IgG (LI-COR, IRDye® 680RD, 926-68070, 1:20,000) secondary antibodies at room temperature for 1 hour. Primary and secondary antibody incubations were done in Intercept (LI-Cor Intercept TBS antibody diluent, 927-65001) buffer plus 0.2% Tween; washes were with TBS plus 0.2% Tween with final rinse in TBS only. After washing, membranes were dried, scanned, and signal was detected using the LI-COR Odyssey CL-X Imaging System and Image Studio software (LI-COR).

| Antibody | Manufacturer | Catalog Number | Dilution |
| --- | --- | --- | --- |
| MSH3 Polyclonal Antibody | Bethyl Labs | A305-314A | 1:1000 |
| GAPDH Monoclonal Antibody (GA1R) | Invitrogen | MA5-15738 | 1:5000 |
| MSH2 Monoclonal Antibody (FE11) | Invitrogen | 33-7900 | 1:1000 |
| Beta-Actin Monoclonal Antibody | Proteintech | 60008-1-1g | 1:5000 |
| Histone H3 Antibody | Cell Signaling Technology | 9715S |  |

### Cell fractionation

CelLytic™ NuCLEAR™ Extraction Kit (Millipore Sigma, NXTRACT-1KT) was used to perform cell fractionation following the manufacturer’s instructions. 2 million cells were harvested and subjected to fractionation. The cell pellets were rinsed twice with PBS buffer. 500 µL (5X PCV) of 1X Lysis Buffer (including 0.01M DTT and protease inhibitors) was added and the cell pellet was resuspended gently. The packed cells were incubated in the selected lysis buffer (isotonic) on ice for 15 minutes, allowing cells to swell. 10% IGEPAL CA-630 solution was added to a final concentration of 0.6%, followed by vortexing vigorously for 10 seconds and immediate centrifugation for 30 seconds at 10,000 x g. The supernatants (cytoplasmic fraction) were transferred to new tubes. The nuclei pellets were washed with isotonic 1x Lysis Buffer twice and then lysed with RIPA buffer on ice for 10 minutes. The lysates were centrifuged for 5 minutes at 11,000 xg and the supernatants were transferred to new tubes.

### Frameshift assay

Flp-In™ 293 cells containing AAAG eGFP reporter constructs but lacking GFP expression were isolated into 6-well plates (400,000 cells per well) at day 0 using a Sony SH800 cell sorter. Parental Flp-In™ 293 cells and AAAG18-GFP (in frame) cells serve as the negative and positive controls respectively. Cells were passaged twice a week at the density of 2.2 ×10^6^ in 10 cm dish. Cytometry analysis was performed at day 7, 14 and 21 using the MacsQuant VYB (Miltenyi Biotec) flow cytometer. Cells were rinsed with cold Ca^2+^/Mg^2+^ -free PBS (GIBCO-Invitrogen), detached with trypsin (ThermoFisher Scientific 25200-056) and neutralized with culture media. After centrifugation (5 minutes at 100 xg), cell pellets were suspended in FACS buffer (PBS with 0.5% BSA, 2mM EDTA, 0.02% sodium azide) to achieve a density of 1 million cells per mL. The MacsQuant flow cytometer was used to acquire approximately 500,000 events. In compound treatment experiments DMSO or RGFP966 were applied to cell cultures at day 1 (24h after sorting). Compound treatment was maintained throughout the course of the assay by including compound in each passage.

Data analysis was performed by FlowJo software. Essentially, forward scatter area (FSC-A) versus side scatter area (SSC-A) gating was used to identify cells of interest based on size and granularity (complexity). Forward scatter height (FSC-H) versus forward scatter area (FSC-A) gating was applied subsequently to identify single cells and ensure doublet discrimination. To quantitate the GFP+ cell population the GFP intensity data was plotted as a histogram. Cell populations that demonstrate instability contain a large intensity peak with background GFP fluorescence and a small secondary peak with higher levels of GFP fluorescence. (In all experiments, the second peak was distinct from the HEK293 parental cell line not containing a GFP reporter construct). The gate for GFP+ cells was drawn at the minimum between the 2 peaks (Figure S3) and the percentage of cells with GFP intensity greater than this value was determined. Percent positive GFP values were plotted vs time, and a line of best fit passing through the origin was calculated using GraphPad Prism. The slope of this line was used to determine the rate of GFP+ conversion per week.

## Author contributions

C.B. and J.F. conceived the concept and initiated the project. J.A., M.A.A., M.M., R.M. and J.F. designed and supervised experiments. T.T., S.C. and J.A. generated and characterized frameshift stable cell lines. J.A., T.T., M.A.A., S.C. and M.M. conducted experiments for collected data. J.A., M.A.A., M.M., R.M., R.N., and J.F. analyzed and interpreted data. J.A. and C.B. drafted the manuscript with input from all authors, R.M., J.F. and C.B. performed critical revision of the manuscript. All authors read, revised, and approved the manuscript.

## Acknowledgments

The authors would like to acknowledge the Pfizer Sequencing Core for the generation of RNA-seq data, and our Pfizer Flowcytometry Center colleagues Matthew Powers and Hanna Sobon for technical support, critical discussion of the fluorescence gating strategy, and assay optimization. We thank Amrutha Pattamatta, Timothy Calamaras, Anthony Razov, and Claudia Huichalaf for critical review and editing of the manuscript.

