## Supplemental Material for "Development of a cellular reporter assay to measure activity of MutSβ, a therapeutic target for Huntington’s disease"

### Methods

**Digital droplet PCR (ddPCR) assay to analyze the expression of known MMR genes in Flp-In™ 293 cells**

To analyze the expression of known MMR genes in FLP293 cells, 20 µl ddPCR reactions were prepared as described previously (Mohiuddin et al., 2022). Briefly, 10 µl of ddPCR™ Supermix for Probes (No dUTP, BioRad, 1863024), 900 nM each of the forward and reverse primers, 250 nM each of the fluorescence probes, 1 µL DNA (60 ng/µL), and RNase/DNase-free water. Two targets were detected simultaneously with different fluorescence probes: the FAM probe detected the specific MMR genes, while the VIC probe detected the HPRT gene. The annealing temperature for different sets of primers and probes was established by previous gradient runs. After droplet generation, PCR was performed in C1000 Touch Thermal Cycler as follows, 95°C 10 min, 40 cycles of 94°C 30s, 58°C 1 min, 98°C 10 minutes. Droplet analysis was performed using the QX200 instrument, and data were analyzed with Quantasoft software, with the absolute quantification (ABS) mode according to the digital MIQE guidelines. Primers are listed in the table below.

| **Target** | **Number** | **Catalog Number (ThermoFisher)** |
|---|---|---|
| PMS1 | 1 | Hs00922261 |
| | 2 | Hs00922262 |
| PMS2 | 1 | Hs01562427 |
| | 2 | Hs00241053 |
| MLH1 | 1 | Hs00979919 |
| | 2 | Hs00179866 |
| MLH3 | 1 | Hs00998142 |
| | 2 | Hs00271778 |
| MSH3 | 1 | Hs00267239 |
| | 2 | Hs00989003 |
| MSH6 | 1 | Hs00264721 |
| | 2 | Hs00943000 |
| MSH2 | 1 | Hs00179887 |

| | | |
|---|---|---|
| HPRT | 1 | Hs02800695 |

**DNA-sequencing**

Standard whole-genome sequencing was performed on the AAAG17 Flp-In™ 293 cells by Fulgent using Illumina Sequencers with 40X coverage. Reads were demultiplexed using Illumina bcl2fastq2 v2.20 with default settings. nf-core/sarek workflow (3.1.2) was run in nextflow (22.10.4) with default settings: quality control and trimming were run with FastQC (0.11.9) and fastp (0.23.2); reads were mapped to the reference GRCh38 genome with BWA-mem (0.7.17-r1188); bam files were processed with GATK (4.3.0.0) MarkDuplicates, BaseRecalibrator, and ApplyBQSR; variant calling was run with GATK (4.3.0.0) HaplotypeCaller. Variants that were in mismatch repair (MMR) genes *MSH3, MSH2, MSH6, MLH1, PMS2, PMS1, MLH3, PCNA, EXO1, FAN1,* and *RPA* were annotated with Ensembl Variant Effect Predictor (VEP) version 101.0, and protein-coding variants were extracted.

### Figure and Table Legends

**S1. Expression of known MMR genes in FLP293 cells.** Expression of MMR genes relative to HPRT were measured by droplet digital PCR (dd-PCR) assay. -1 and -2 designations indicate the use of 2 separate primer/probe sets used for detection.

**S2. Determining number of integration events in reporter positive clones.** Droplet digital PCR results showing the number of copies of the GFP reporter relative to hTERT in the genome. The expected ratio for a single integration event is 0.33 since 293 cells have trisomy of chromosome 5.

**S3. Example flow data illustrating gating strategy.** Dot plot (left) and histogram (right) of GFP signal in example flow data.  GFP positive cells were defined as the distinct second population of cells observed in the histogram plots. Cell with in-frame reporter construct and cells without integration of reporter construct were used to corroborate gates.

**S4. Experimental reproducibility with $AAAG_{17}$ reporter.** Intraexperimental (left), interexperiment (middle) and clonal variability (right) in the rates of GFP+ conversion in $AAAG_{17}$ lines are shown.

**S5. Repeat number distribution in GFP negative cells.** Distribution of repeat length in RNAseq reads from GFP- subpopulation of $AAAG_{17}$ reporter cell lines. Green color represents repeat numbers that would produce in frame GFP protein expression.

**S6. Evaluation of $AAAG_{31}$ reporter lines.** A. GFP+ conversion in $AAAG_{31}$ reporter lines expressing shRNA directed toward MSH3. B. GFP+ conversion in $AAAG_{31}$ reporter MSH3 knockout lines with and without lentiviral mediated expression of wild-type MSH3. C. Western blot probing MSH3 protein levels in $AAAG_{31}$ wild-type cells and cells stably expressing shRNA directed toward MSH3 (left panel), and in MSH3 knockout cells complemented with WT MSH3 (right panel).

**Figure S7. Extent of GFP fluorescence with varying AAAG repeat length.** Dot plots of the log of GFP signal intensity vs. side scatter from the indicated AAAG reporter strains.

**Figure S8. MSH3 expression in $AAAG_{13}$ complemented cells.** Western blot probing MSH3 protein levels in $AAAG_{13}$ wild-type cells, MSH3 knockout cells, and MSH3 knockout cells complemented with either WT or E976A MSH3.

**Figure S9. Evaluation of coding polymorphisms in human genome that modulate disease parameters in HD.** A. Alignment of coding polymorphisms in N terminus of MSH3. Allele 3a has a deletion which removes 9 amino acids in the N terminal repeat region relative to the major allele (6a). Polymorphism 7a contain an insertion that results in the introduction of 3 amino acids relative to the major allele. B. Kinetics of GFP+ conversion in wild-type (dark blue), MSH3 KO (red), and MSH3 KO cells expessing the major allele (6a – green), or the minor alleles 3a (purple) or 7a (orange). Error bars represent standard deviation (n=2). C. Western blot probing MSH3 protein levels in $AAAG_{17}$ wild-type cells, MSH3 knockout cells, and MSH KO cells expressing the indicated MSH3 allele.

**Table S1: DNA variants in MMR genes in Flp-In™ 293 cells.** Only protein-coding variants are shown. Genomic positions are in GRCh38.

**Table S2: $AAAG_{17}$ RNA-seq repeat counts.** The percentage of RNA-seq reads with various repeat counts across the reporter repeat region in the given cell populations. Fractional repeat counts represent potential insertions/deletions of < 4 nucleotides in the repeat region.

**Table S3. $AAAG_{13}$ RNA-seq repeat counts.** The percentage of RNA-seq reads with various repeat counts across the reporter repeat region in the given cell populations. Fractional repeat counts represent potential insertions/deletions of < 4 nucleotides in the repeat region.
