## Supplemental Figures for "Development of a cellular reporter assay to measure activity of MutSβ, a therapeutic target for Huntington’s disease"

### Supplemental Figure 1

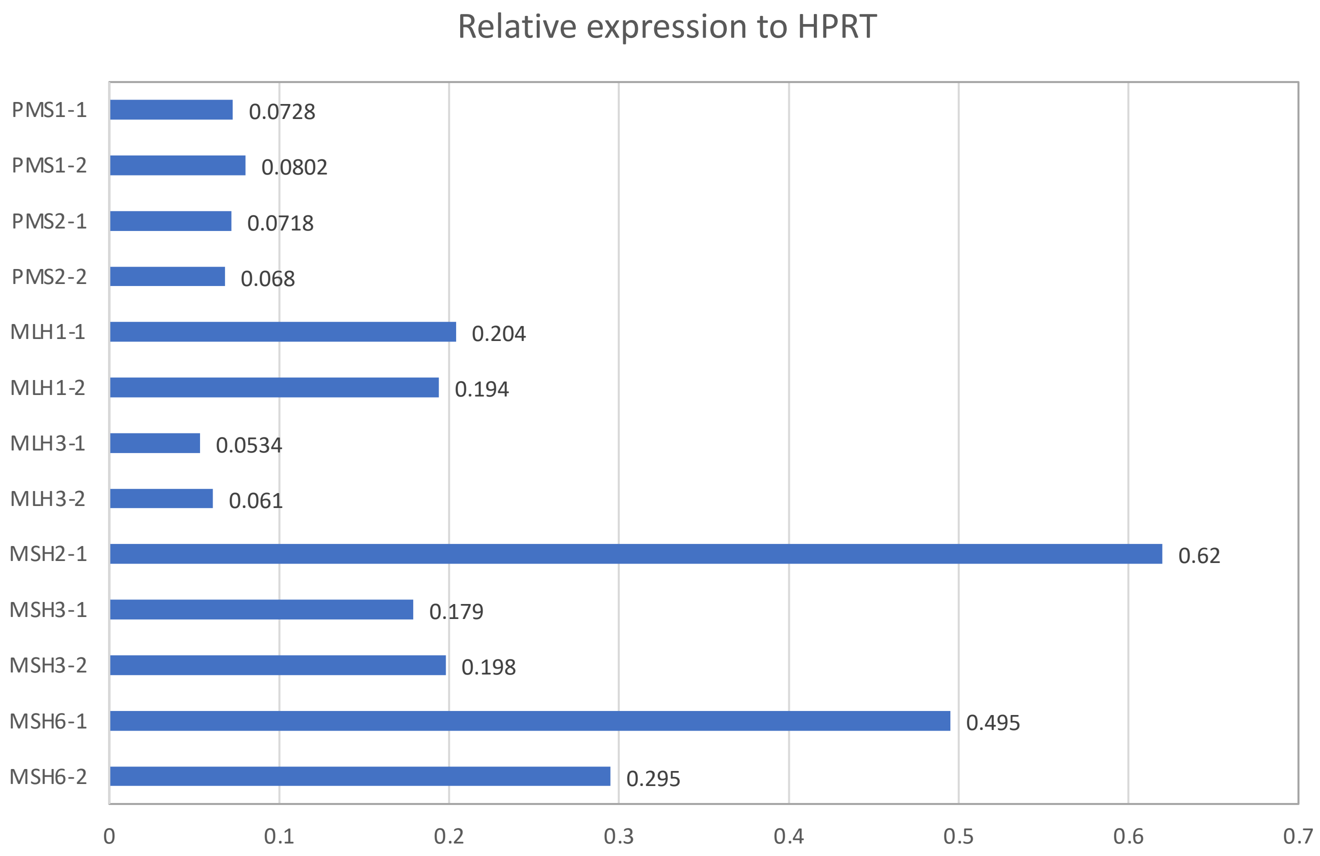

### Supplemental Figure 2

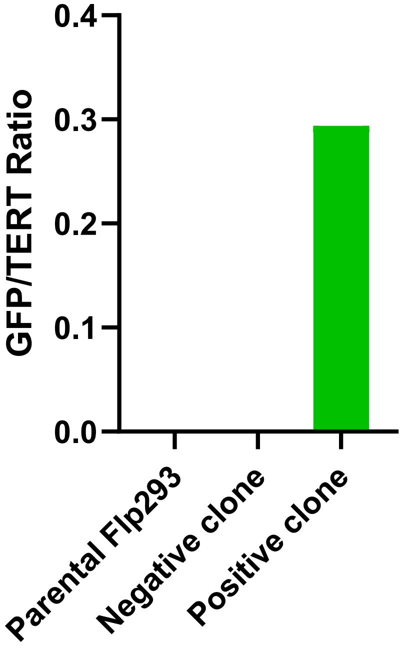

### Supplemental Figure 3

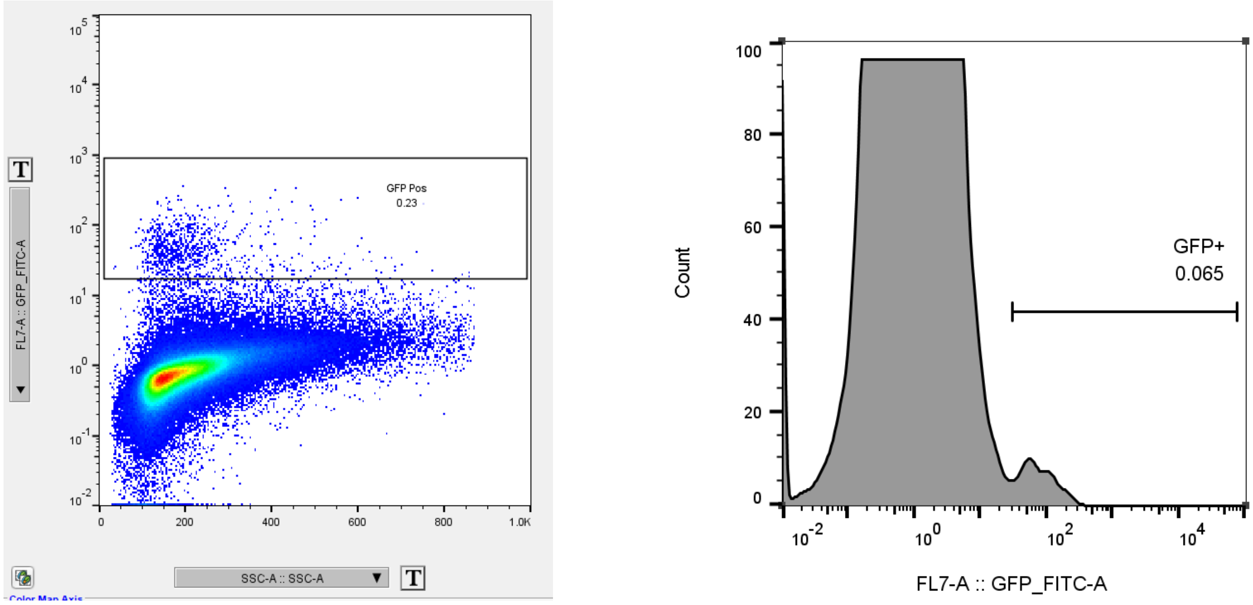

### Supplemental Figure 4

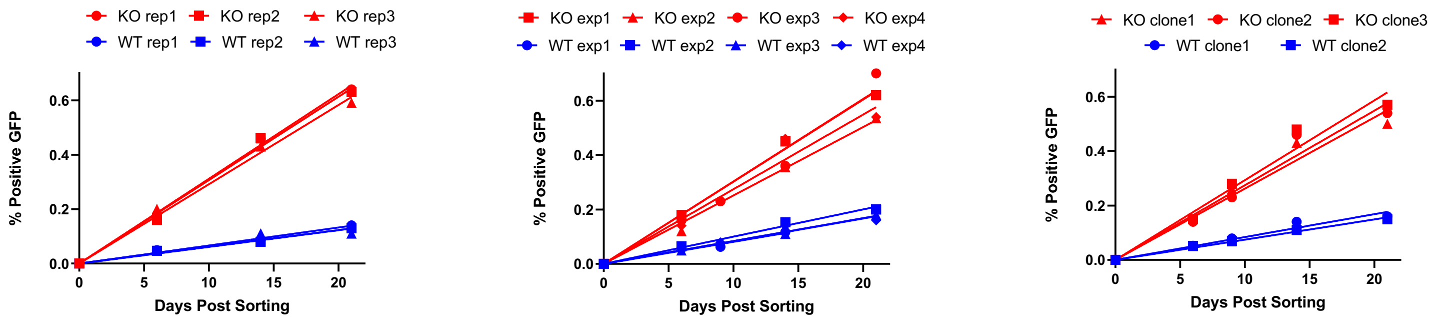

### Supplemental Figure 5

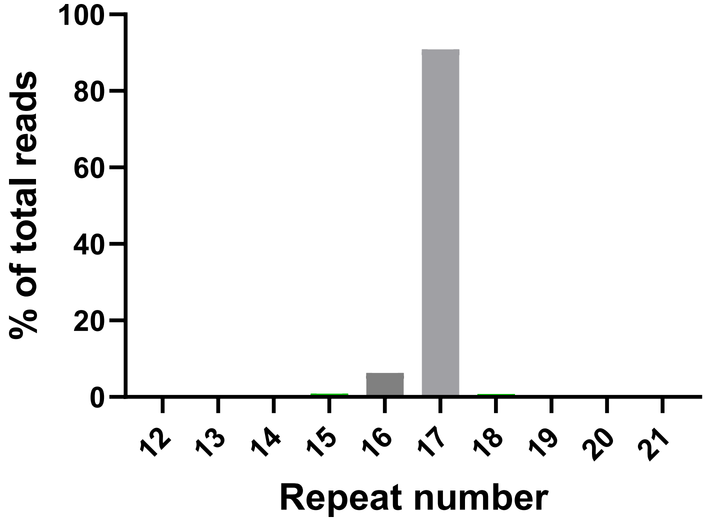

### Supplemental Figure 6

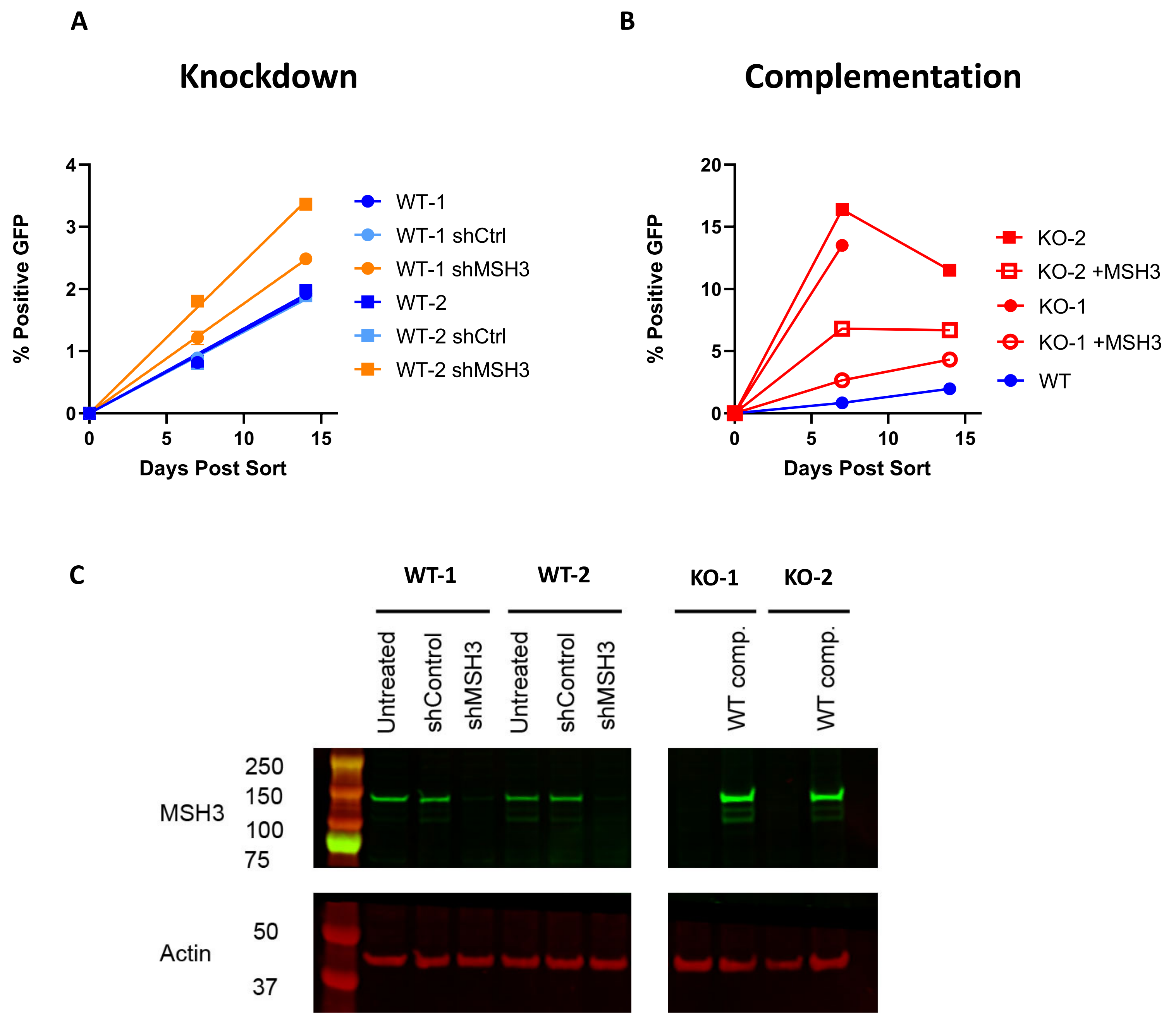

**Supplemental Figure 7**

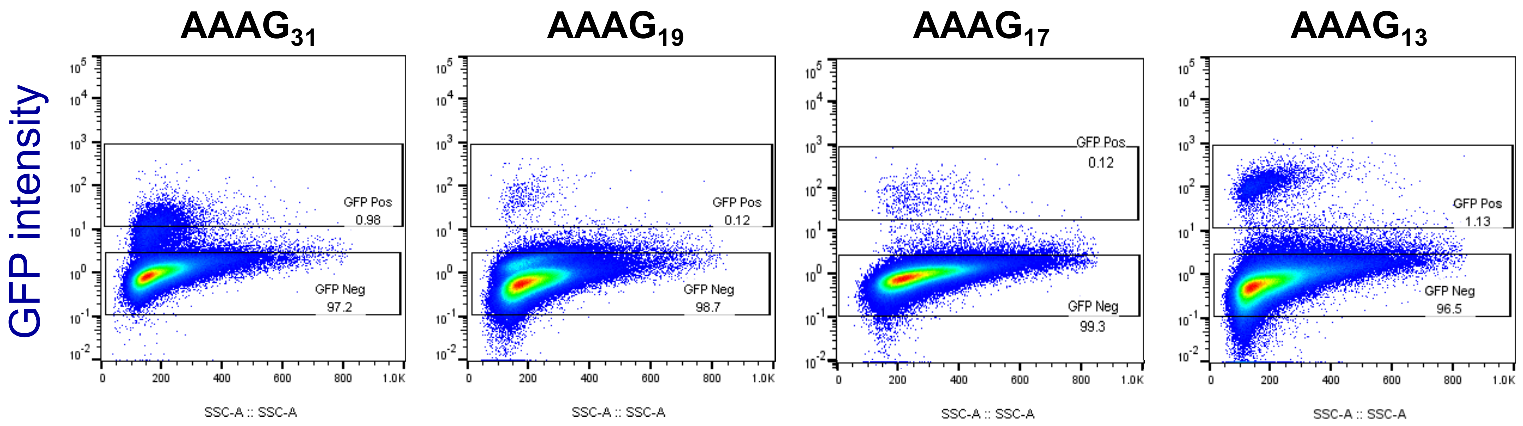

### Supplemental Figure 8

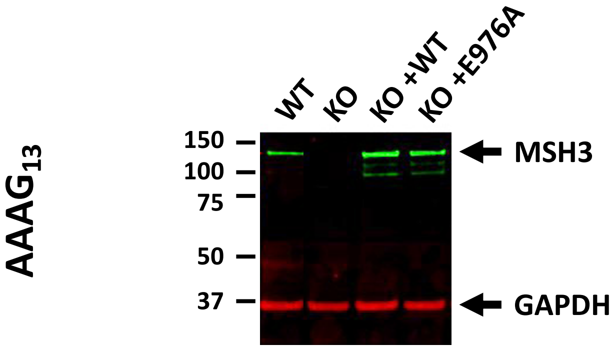

### Supplemental Figure 9

**A**

```
6a  (Major Allele)     45  DQVDPGAAAAAAAAAA---AAPPAPPAPAFPPQLPPH  78
3a                         DQVDPGAAAAAA------------APPAPAFPPQLPPH
7a                         DQVDPGAAAAAAAAAAAAPAAPPAPPAPAFPPQLPPH
```

**B**

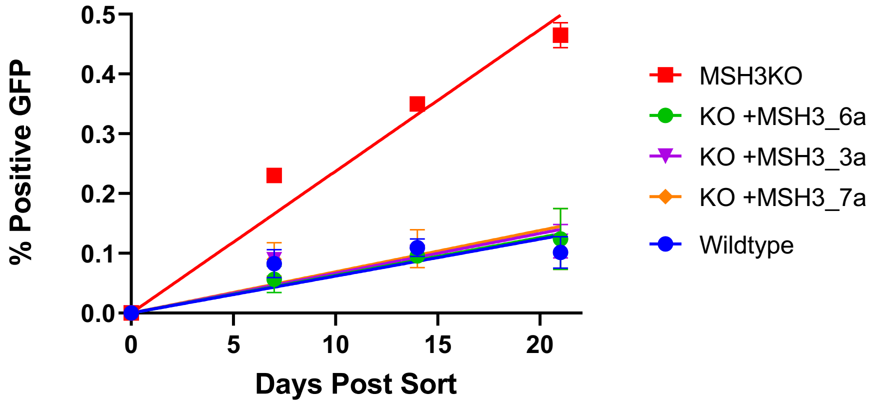


**C**

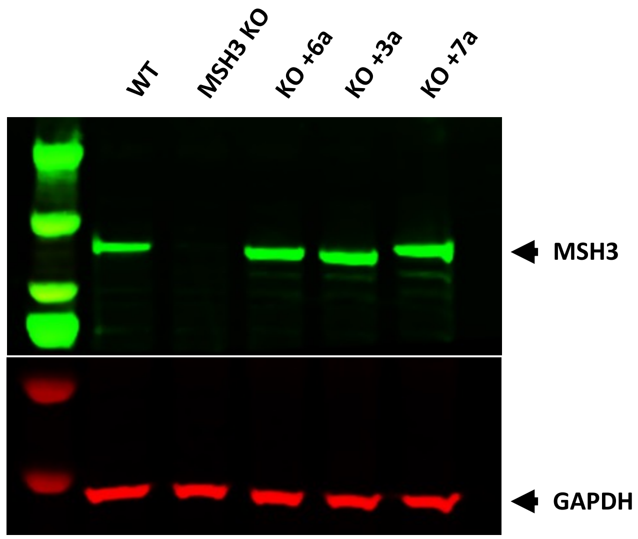

### Supplemental Table 1

| Gene | Chrom | Position | Ref | Alt | Protein Change | Zygosity | Variant type | gnomAD v3.1.2 allele frequency | ClinVar annotation |
|---|---|---|---|---|---|---|---|---|---|
| EXO1 | chr1 | 241866849 | A | G | p.His354Arg | het | missense | 0.593 | Not in ClinVar |
| EXO1 | chr1 | 241878999 | G | A | p.Glu589Lys | hom | missense | 0.399 | Not in ClinVar |
| EXO1 | chr1 | 241881973 | C | T | p.Arg723Cys | hom | missense | 0.9459 | Not in ClinVar |
| FAN1 | chr15 | 30905361 | G | A | p.Gly233Glu | hom | missense | 0.3669 | Benign |
| FAN1 | chr15 | 30937217 | T | C | p.His1005= | hom | synonymous | 0.3937 | Benign |
| MLH3 | chr14 | 75047180 | T | C | p.Asn826Asp | hom | missense | 0.989 | Benign |
| MLH3 | chr14 | 75048398 | C | T | p.Val420Ile | het | missense | 0.01089 | Benign/Likely benign |
| MLH3 | chr14 | 75017109 | T | C | p.Gln1445= | hom | synonymous | 0.5262 | Benign |
| MSH2 | chr2 | 47512412 | A | G | p.Gln915Arg | het | missense | 0.4626 | Benign |
| MSH3 | chr5 | 80854162 | A | G | p.Gln949Arg | hom | missense | 0.8671 | Benign |
| MSH3 | chr5 | 80873118 | G | A | p.Ala1045Thr | hom | missense | 0.706 | Benign |
| MSH6 | chr2 | 47795976 | T | C | p.Asp180= | het | synonymous | 0.2314 | Benign |
| MSH6 | chr2 | 47798625 | C | T | p.Tyr214= | het | synonymous | 0.07188 | Benign |
| PCNA | chr20 | 5115453 | G | C | p.Pro234= | het | synonymous | 0.001623 | Benign |
| PMS2 | chr7 | 5973418 | C | G | p.Gly857Ala | het | missense | 0.3103 | Benign |
| PMS2 | chr7 | 5987144 | T | C | p.Lys541Glu | hom | missense | 0.8654 | Benign |
| PMS2 | chr7 | 5987357 | G | A | p.Pro470Ser | hom | missense | 0.372 | Benign |
| PMS2 | chr7 | 5997349 | G | C | p.Ser260= | hom | synonymous | 0.8169 | Benign |

### Supplemental Table 2

| Repeat length | Repeat count | % of reads | | | |
|---|---|---|---|---|---|
| | | Unsorted WT | WT GFP+ | WT GFP- | KO GFP+ |
| 24 | 6 | 0.000% | 3.212% | 0.000% | 0.000% |
| 28 | 7 | 0.000% | 0.079% | 0.020% | 0.071% |
| 32 | 8 | 0.132% | 0.079% | 0.000% | 0.142% |
| 36 | 9 | 0.000% | 2.463% | 0.041% | 0.283% |
| 40 | 10 | 0.000% | 0.158% | 0.031% | 0.142% |
| 42 | 10.5 | 0.000% | 0.059% | 0.000% | 0.000% |
| 44 | 11 | 0.066% | 0.650% | 0.000% | 0.389% |
| 48 | 12 | 0.397% | 8.571% | 0.163% | 3.892% |
| 52 | 13 | 0.066% | 1.419% | 0.132% | 0.672% |
| 53 | 13.25 | 0.000% | 0.020% | 0.000% | 0.000% |
| 54 | 13.5 | 0.000% | 0.138% | 0.000% | 0.000% |
| 56 | 14 | 0.661% | 4.433% | 0.437% | 4.069% |
| 57 | 14.25 | 0.000% | 0.926% | 0.000% | 0.000% |
| 59 | 14.75 | 0.000% | 0.039% | 0.000% | 0.000% |
| 60 | 15 | 1.718% | 44.099% | 0.834% | 47.629% |
| 63 | 15.75 | 0.000% | 0.039% | 0.041% | 0.000% |
| 64 | 16 | 6.742% | 2.365% | 6.256% | 3.114% |
| 66 | 16.5 | 0.000% | 0.000% | 0.010% | 0.000% |
| 67 | 16.75 | 0.529% | 0.020% | 0.275% | 0.000% |
| 68 | 17 | 86.319% | 4.473% | 90.886% | 6.192% |
| 69 | 17.25 | 0.066% | 0.000% | 0.020% | 0.000% |
| 70 | 17.5 | 0.000% | 0.000% | 0.000% | 0.106% |
| 71 | 17.75 | 0.000% | 0.000% | 0.000% | 0.071% |
| 72 | 18 | 2.776% | 24.650% | 0.793% | 31.458% |
| 76 | 19 | 0.397% | 1.478% | 0.031% | 1.380% |
| 80 | 20 | 0.132% | 0.433% | 0.031% | 0.248% |
| 84 | 21 | 0.000% | 0.138% | 0.000% | 0.142% |
| 88 | 22 | 0.000% | 0.039% | 0.000% | 0.000% |
| 104 | 26 | 0.000% | 0.020% | 0.000% | 0.000% |

### Supplemental Table 3

| Repeat length | Repeat count | % of reads | |
|---|---|---|---|
| | | Unsorted | GFP+ |
| 24 | 6 | 0.559% | 0.086% |
| 28 | 7 | 0.140% | 0.000% |
| 32 | 8 | 0.279% | 0.301% |
| 35 | 8.75 | 0.000% | 0.086% |
| 36 | 9 | 0.140% | 15.606% |
| 40 | 10 | 0.140% | 1.032% |
| 44 | 11 | 2.933% | 4.213% |
| 48 | 12 | 4.888% | 76.526% |
| 52 | 13 | 87.151% | 2.021% |
| 56 | 14 | 3.771% | 0.086% |
| 60 | 15 | 0.000% | 0.043% |
